# Adaptive Unscented Kalman Filter for Neuronal State and Parameter Estimation

**DOI:** 10.1101/2022.06.29.497821

**Authors:** Loïc J. Azzalini, David Crompton, Gabriele M. T. D’Eleuterio, Frances Skinner, Milad Lankarany

**Affiliations:** Institute for Aerospace Studies, University of Toronto, Toronto, Ontario, Canada; Division of Clinical and Computational Neuroscience, Krembil Research Institute, University Health Network, Toronto, Ontario, Canada; Department of Medicine (Neurology), University of Toronto, Toronto, Ontario, Canada; Department of Physiology, University of Toronto, Toronto, Ontario, Canada; Institute of Biomedical Engineering, University of Toronto, Toronto, Ontario, Canada

**Keywords:** Conductance-based model, Nonlinear Kalman Filtering, Adaptability, Model Mismatch

## Abstract

Data assimilation techniques for state and parameter estimation are frequently applied in the context of computational neuroscience. In this work, we show how an adaptive variant of the unscented Kalman filter (UKF) performs on the tracking of a conductance-based neuron model. Unlike standard recursive filter implementations, the robust adaptive unscented Kalman filter (RAUKF) jointly estimates the states and parameters of the neuronal model while adjusting noise covariance matrices online based on innovation and residual information. We benchmark the adaptive filter’s performance against existing nonlinear Kalman filters and explore the sensitivity of the filter parameters to the system being modelled. To evaluate the robustness of the proposed solution, we simulate practical settings that challenge tracking performance, such as a model mismatch and measurement faults. Compared to standard variants of the Kalman filter the adaptive variant implemented here is more accurate and robust to faults.

## 1 Introduction

The application of data assimilation (or *state estimation*) techniques to single neuron dynamics was greatly popularized by Schiff (2011), based on the work of Voss, Timmer, and Kurths (2004) on the FitzHugh-Nagumo model. The latter have shown that recursive Bayesian state estimators such as the unscented Kalman filter (UKF) (Julier & Uhlmann, 1997) could be used to track the nonlinear dynamics of neuronal models and identify relevant model parameters based on the observation of a measurable, albeit noisy, membrane voltage trace. The effective combination of tracking and system identification garnered a lot of interest from researchers working at the intersection of computational neuroscience and control theory. While indispensable for attitude estimation in aerospace and localization in robotics (Barfoot, 2017), state estimation is still emerging in the neuroscience and biomedical fields.

While often assumed to be static, changes in neuron model parameters can lead to significantly different excitability characteristics. In the absence of direct measurements of such parameters, recursive state estimators allow for observed states, unobserved states, and parameters to be tracked more accurately. Moye and Diekman (2018) explore the robustness of data assimilation against poor initialization of neuronal parameters by comparing the performance of the UKF, a sequential approach, to variational methods. The UKF has been shown to be robust against mismatches between the model known *a priori* and the model from which the observed data originates, even in the presence of significant model inaccuracies (e.g., steady-state constants replacing transient dynamics (Ullah & Schiff, 2009)). As knowledge of these underlying models may be lacking (particularly in neuroscience), adaptive techniques that simultaneously identify missing parameters are highly desirable. This challenge is common to many disciplines and it has led to the concept of robust adaptive unscented Kalman filter (RAUKF) (Hajiyev & Soken, 2014; Zheng, Fu, Li, & Yuan, 2018). However, to the best of our knowledge, these robust and adaptive methods have yet to be applied to neuronal dynamics.

The Kalman filter is known to be the optimal recursive state estimator in the context of linear dynamics and Gaussian distributed noise (Kalman, 1960). However, neuronal dynamics are typically highly nonlinear and warrant the use of nonlinear estimators such as the extended Kalman filter (EKF), unscented Kalman filter (UKF) (Julier, Uhlmann, & Durrant-Whyte, 2000) or particle filter (PF). While linearization of neuronal dynamics about the most recent state estimate (such as in the EKF) can be shown to perform well in certain situations, it is prone to instability and divergence (Lankarany, Zhu, & Swamy, 2014). Deterministic sampling alternatives, such as the sigma-point transform around which the UKF is built, are generally more suited to the dynamics under study here (Schiff, 2011). In this case, the analytical derivatives of the dynamics and observation models used in the EKF are no longer required. Instead, the sampling-based UKF allows for the models to be treated as black-boxes, which could suit practical biomedical applications.

Despite the undoubted capability of these techniques in inferring hidden dynamics in nonlinear systems, some critical challenges related to the robustness of UKF and other families of KFs remain when it comes to their application on real time inference on biological models. These challenges are mainly related to a lack of *a priori* information to inform the initialization of state variables and noise covariance matrices. When modelling neurons, one must take into account the behaviour of specific ion channels, their conductances, and kinetics, which may not be present in the dataset to provide suitable estimates for the initial state, especially as conductances and dynamics vary greatly in different neurons, even within the same region (Golowasch, 2014). The inability of the model to observe the full state, as well as abrupt changes that can occur in recordings (e.g., sharp discontinuities in membrane potential traces) can lead to drastic changes in covariance estimates and in turn, instability. While ad hoc adjustments of covariance matrices such as covariance inflation have been proposed in the past (Schiff, 2009), a state estimation method capable of handling biological systems is still lacking. The present work endeavours to address this gap.

To address the challenges of applying the KF to neuron models, we consider a few modifications based on modern adaptations of the UKF. For KF initialization, we address this challenge as follows: We employ a RAUKF which, through online fault detection, adjusts the covariances supplied to more appropriate values. We ensure optimal performance of this fault detection by performing a grid space search of the relevant parameters across a range of trials. The optimal parameters determined are model specific; however, certain aspects of the values determined may be used with some consideration of their relevance in what parameters need to be estimated. For the concern of the model being incomplete and thus unable to represent fully or reproduce the desired behaviour, we implement a more detailed model and track it via RAUKF using the less complete version of the model. In doing so, we monitor the performance over time, especially when the less complete model is unable to match spike times, hyperpolarization curves, or other features due to its incompleteness. Abrupt changes in recorded data, be they from random noise or artifacts from the recording, are addressed by the robust and adaptive response to state changes in the RAUKF.

In this paper, we develop a RAUKF for neuronal state and parameter estimation based on the work of Hajiyev and Soken (2014) and Zheng et al. (2018). We use two variants of the conductance-based Morris-Lecar neuron model (Prescott, De Koninck, & Sejnowski, 2008) to showcase the filter’s adaptability against noise and lacking model information.

The following section (2.1) introduces the 2-dimensional conductance-based model, which is subsequently used in the state estimation framework (2.2) as the dynamics model. The core implementation of the UKF is reviewed in section 2.3 before introducing the extensions that support adaptation (2.4) and correction (2.5) of uncertain parameters and noise covariance matrices.

A numerical exploration of the RAUKF parameter space is provided in section 3.1 to identify the set of conditions best suited for this application. The performance of the RAUKF for neuronal state estimation is compared to existing algorithms in section 3.2, followed by simulations mimicking measurement faults (3.3) and model discrepancies (3.4). The significance of the results for neuronal dynamics identification and future extensions are discussed in section 4.

Overall, the RAUKF outperforms a standard UKF implementation in the case of poor initialization of covariance matrices and neuronal model parameters. The bulk of RAUKF adaptation steps are taken at the onset of simulation, where it adjusts its parameters based on the most recently observed data. The advantages of adaptation are particularly noticeable in the presence of measurement faults and model mismatch where the performance of a standard implementation quickly deteriorates. Demonstrations of robustness against experimental faults are of particular importance to validate the use of algorithms like RAUKF in practical settings.

## 2 Method

### 2.1 Neuron Model

As the simplest possible biophysical representation for a neuron, conductance-based models are commonly used in computational neuroscience (Skinner, 2006). To demonstrate applications of this filtering technique, we use variants of the Morris-Lecar model as provided by Prescott et al. (2008) (see Appendix A). As our developed method may be applied to any conductance-based neuron model, we here refer to a generic conductance-based neuron model as the neuron model:

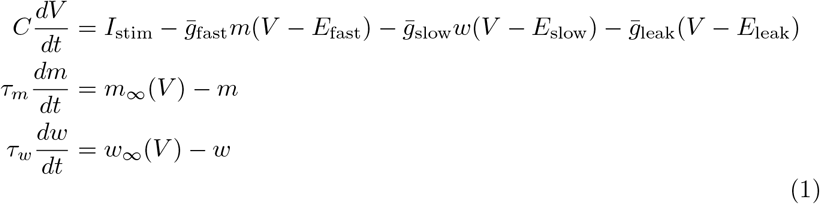

where *V* denotes the membrane voltage, *m* and *w* are arbitrary gating variables with associated time constants *τ_m_* and *τ_w_*, *ḡ*_*_ are maximal conductances and *E*_*_ are ion channel reversal potentials. By considering a separation of timescales, the quasi-steady-state approximation *m* = *m*_∞_ will be used in the following sections. Table 1 provides ranges for each variable.

**Table 1.** Parameter range estimates

| Parameter | Range | Units |
| --- | --- | --- |
| $V$ | $[-100, 50)$ | mV |
| $w$ | $[0, 1]$ | |
| $m$ | $[0, 1]$ | |
| $\bar{g}_{\text{fast}}$ | $[0.1, 100]$ | $\text{mS cm}^{-2}$ |
| $\bar{g}_{\text{slow}}$ | $[0.1, 100]$ | $\text{mS cm}^{-2}$ |
| $\bar{g}_{\text{leak}}$ | $[0.1, 10]$ | $\text{mS cm}^{-2}$ |

### 2.2 Neuronal State and Parameter Estimation

In this context, the main goal of data assimilation consists in estimating *V* and *w* based on noisy measurements of the membrane voltage and predictions 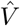 and *ŵ* afforded by the dynamics model (1). Given that the incoming measurements represent the newest source of information about the biological system, the recursive updates are performed at a rate equal to the measurement sampling period *T*. To accommodate this discrete process, the dynamics are discretized, with state **x**_*k*_ = [*V_k_ w_k_*]^*T*^ and input current **u**_*k*_ = *I_stim,k_*:

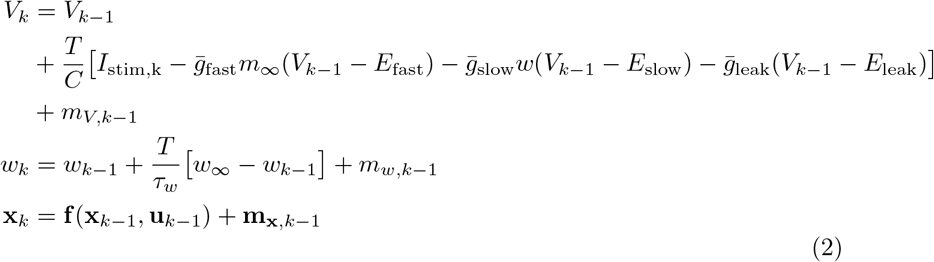

where *k* = 1,…, *N*, 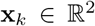, 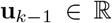 and 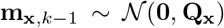 models uncertainty inherent to the dynamics.

Given a neuron model such as (1), attributing the notion of internal state to the membrane voltage *V* and recovery variable *w* to comply with the statespace representation of control theory (2) can be ambiguous. As alluded to in section 1, the geometrical space of a model describing a biological system relies on a set of parameters which are not constant in reality. The parameters reflect biophysical processes that fluctuate as a result of noisy interactions. While we can initialize parameters based on ranges obtained from experimental studies (see Table 1), parameter estimates more consistent with the latest observations are desired. The conductance of the ionic channels in (1) are a prime example of biologically relevant parameters that naturally vary at a much slower rate of change than the state variables. Consequently, it may be beneficial to directly estimate these parameters from data alongside the state **x**_*k*_.

In joint state and parameter estimation, the state variable is augmented to account for *l* parameters 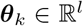 of interest (Stengel, 1994). It is assumed that the rate of change of the parameters is much slower than that of the main variables **x**_*k*_. As such, the parameters are assigned artificial stochastic dynamics

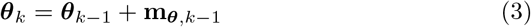

where
**m**_***θ***, *k*–1_ denotes additive white Gaussian noise. The dynamics of the augmented state **X**_*k*_ = [**x**_*k*_ **θ**_*k*_]^*T*^ can then be expressed as

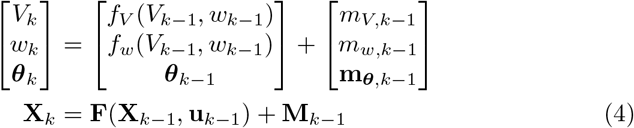

where 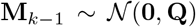. The observation model used to characterize noisy membrane voltage measurements is described by

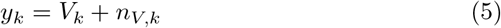

where *n_V,k_* denotes measurement noise (in the general case, 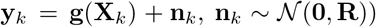. Here, the direct observation of the membrane voltage *y_k_* mimics experimental recording techniques (e.g., current-clamp methods). With only a subset of **X**_*k*_ being measurable, the method presented in this study allows hidden states to be estimated.

### 2.3 Unscented Kalman Filter

Following a Bayesian inference approach and assuming Gaussian beliefs, the state 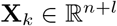 of the neuronal system is a random variable tracking the mean of a normal probability density function, while a covariance 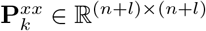 tracks its spread. State estimation consists in estimating the current state **X**_*k*_ given knowledge of its initial conditions **X**_0_, inputs **u**_*k*–1_ and observations of its behaviour over time **y**_*k*_. The filter aims to maximize the probability of observing **y**_*k*_ given a belief about **X**_*k*_ afforded by a model of the system dynamics and an observation model, (4) and (5) respectively. This relationship can be reversed via Bayes’ rule to solve for the *a posteriori* conditional distribution *p*(**X**_*k*_ | **y**_*k*_, **u**_*k*–1_) which represents the true state probability given measurements.

We start by assuming the following Gaussian priors for the prediction step:

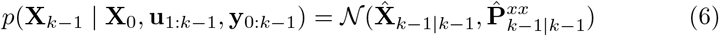

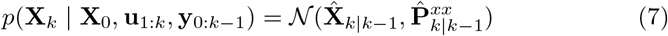

where 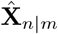 denotes an estimate of **X** at time *n* which includes observations up to and including time *m* ≤ *n* with corresponding estimated covariance matrix 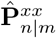. Then, the predicted belief 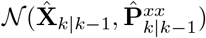 is approximated as follows:

1. A set of 2*N* + 1 sigma-points 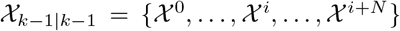 is sampled from the prior density 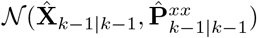 according to

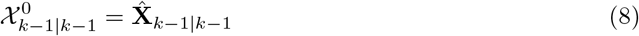

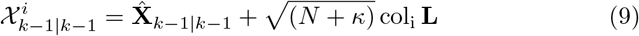

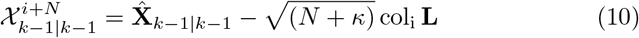

where col_i_ **L** is the *i*th column of the matrix obtained by Cholesky decomposition 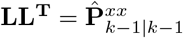; *κ* is a user-definable parameter, often selected according to the heuristic *N* + *κ* = 3 to best capture higher order moments (Julier et al., 2000).
2. The set of sigma-points 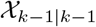 is passed through the nonlinear dynamics model (4)

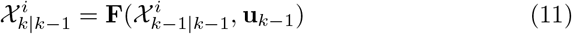
3. The transformed sigma-points are combined into the predicted mean 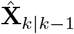 and predicted covariance 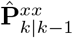

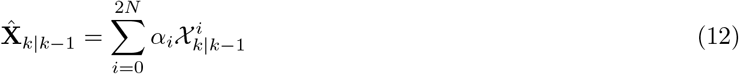

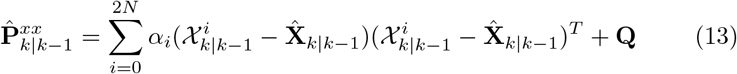

with the weights *α_i_* summing to 1 according to

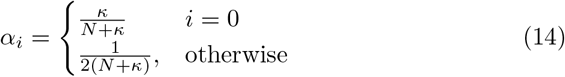

Second, the predicted belief is revised against the most recent observations **y**_*k*_ according to the following steps:

1. A new (optional) set of 2*N* + 1 sigma-points 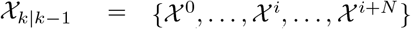 is sampled from the predicted belief 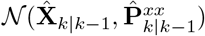 following the same procedure as before
2. The sigma-points 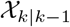 is passed through the measurement model (5)

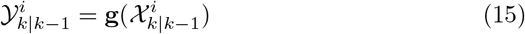
3. The innovation vector **v**_*k*|*k*–1_ and associated covariance matrix 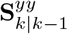 are defined as

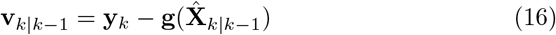

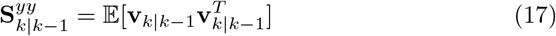
4. The transformed sigma-points are combined into the predicted innovation covariance matrix 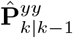 and cross-covariance matrix 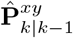

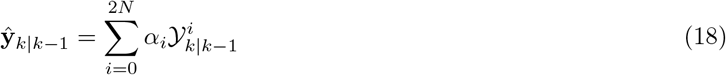

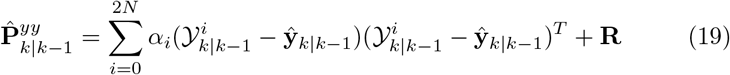

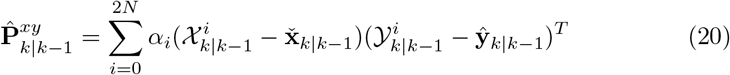
5. The calculation of the Kalman gain **K**_*k*_ and the corrected belief 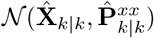 follow from the standard equations below (Barfoot, 2017):

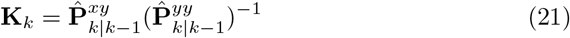

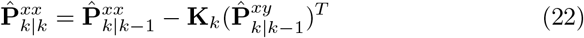

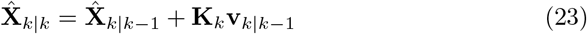
6. Finally, the residual vector **v**_*k*|*k*_ and associated covariance matrix 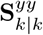 are defined as

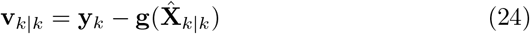

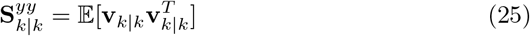
7. Following the same procedure as before, a set of 2*N* + 1 sigmapoints 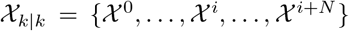 is sampled from the corrected belief 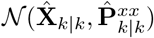, passed through the measurement model (5) and recombined into the predicted residual covariance matrix 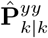

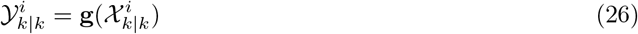

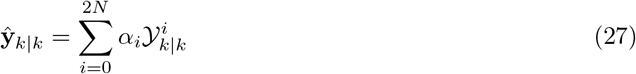

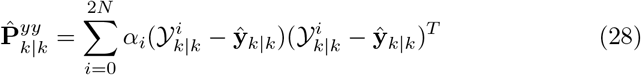

### 2.4 Adaptive Filter

Despite having no guarantee of convergence, the UKF performs well with nonlinear systems, both in tracking state variables and in identifying system parameters. A strong condition for successful estimation is the initialization of covariance matrices **Q** and **R** based on *a priori* information about the system (e.g., measurement noise can be estimated from preexisting data and sensor characteristics). However, in many real-time scenarios, particularly in neuroscience, such information might be lacking, resulting in incomplete initial estimates and in turn, suboptimal filtering (Stengel, 1994). With **Q** and **R** too large, the solution might end up biased, too small and divergence could occur (this is particularly true for slow states, given that they depend on the noise evolution defined by the process noise covariance).

Adaptive techniques have been devised to render the state estimation algorithm more robust against poor estimates of covariance matrices (Schiff, 2009). While methods such as covariance inflation aim to adjust the covariance of the state, and thus improve the stability of the filter, few works have looked at addressing inaccurate noise covariance matrices in this field. Yet, innovation and residual-based approaches (Stengel, 1994) allow, respectively, **Q**_*k*–1_ and **R**_*k*_ to be updated recursively alongside states and parameters based on information readily available in the current UKF implementation.

From (2), (12) and (23),

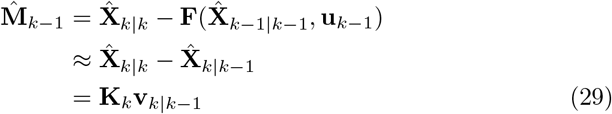

Then,

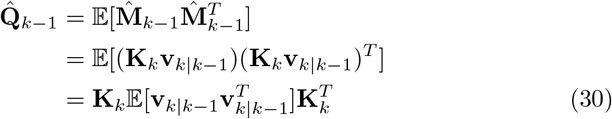

Windowing methods are typically used to approximate 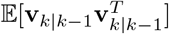 as a sample covariance (Mohamed & Schwarz, 1999; Stengel, 1994)

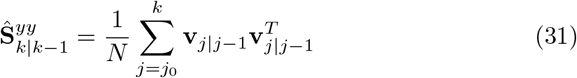

where *j*_0_ = *k* – *N* +1. Alternatively, (Zheng et al., 2018) proposed a weighted update rule based on the direct approximation of the covariance, 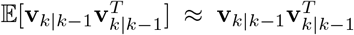 (*N* = 1, effectively). At each adjustment, 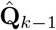 and **Q**_*k*–1_ are combined based on a weighting factor *λ* ∈ (0,1):

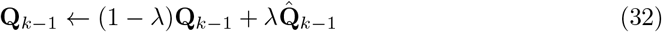

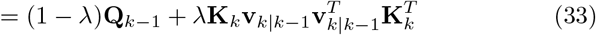

A similar approach is used to estimate 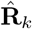, this time based on the residual **V**_*k*|*k*_. From (5),

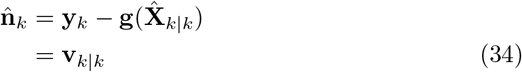

Then,

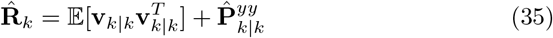

where, similar to (19), 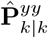 is the covariance of the residual **V**_*k*|*k*_ approximated by the set of 2*N* + 1 sigma-points 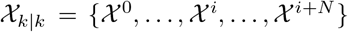 sampled from the corrected belief 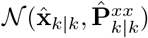 following the same procedure as before.

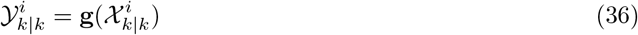

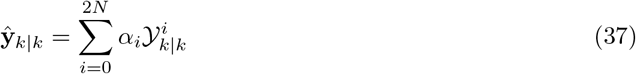

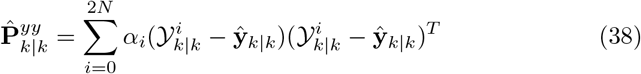

Again, 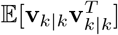 can be approximated by the sample covariance 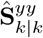 (based on *N* samples) or by a weighted update rule based on a single sample:

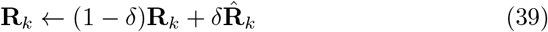

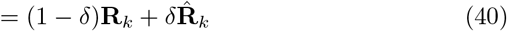

where *δ* ∈ (0, 1) is the weighting factor analogous to *λ*.

### 2.5 Fault Detection

Naturally, the addition of adaptation constitutes a trade-off between computational cost and tracking accuracy. Once sufficiently corrected, additional updates of the noise covariance matrices **Q**_*k*–1_ and **R**_*k*_ will lead to marginal improvements in performance. For this reason, adaptation may be considered as a response to fault detection. Provided with a means of identifying faults in the system (process or measurement-related), adaptation may be used selectively to return the estimation system into normal operating conditions.

A simple fault detection rule follows from innovation-based methods and the statistical function

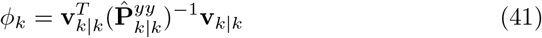

which has *χ*^2^ distribution with *s* =1 degrees of freedom since 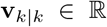 (Hajiyev & Caliskan, 2003; Zheng et al., 2018). Under normal operating conditions, the innovation vector is normally distributed 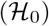. Any deviation from this nominal behaviour could indicate a system fault, such as a damaged sensor, and trigger a recovery mechanism as a result 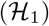. To determine which hypothesis should be accepted, a chi-squared test is performed to determine when 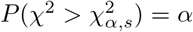, where *α* is the significance level and 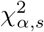 denotes the threshold to be exceeded for a fault to be detected (Hajiyev & Soken, 2014):

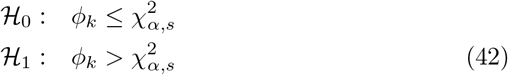

The updates (32) and (39) are performed as a result of rejecting 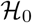. If a windowing method is followed, the selection of *N* effectively tunes the detection system’s sensitivity to faults (a large N may smooth the effects of a potential fault, whereas a small N may lead to false alarms) (Hajiyev & Caliskan, 2003). Alternatively, Zheng et al. (2018) show how the fault threshold could be used in the selection of appropriate weighting factors *λ* and *δ*. Application-dependent tuning parameters *a* > 0 and *b* > 0 are introduced to control how sensitive the noise covariance update rules should be to the innovation statistic *φ_k_*.

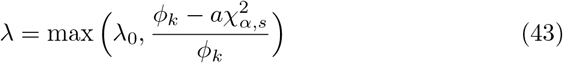

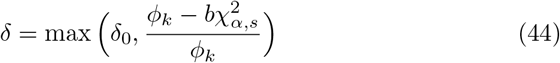

where *λ*_0_ ∈ (0, 1) and *δ*_0_ ∈ (0, 1) are now the default weighting parameters.

## 3 Results

Given the dynamics model (2) and observation model (5), the state **x**_*k*_ may be recursively estimated for *k* = 1,…, *N* using the RAUKF algorithm despite significant amounts of noise in the measurements, poor parameter initialization as well as poor initial estimates of **Q**_0_ and **R**_1_. The noisy measurements *y*_1:*N*,meas_ are obtained beforehand by evolving (2), sampling the membrane voltage at a period *T* = 0.1 ms and adding white Gaussian noise with covariance cov(n_V_) = 3 mV. Figure 1 illustrates these measurements as well as the noisy input stimulation, generated according to an Ornstein-Uhlenbeck process with time constant *τ*_noise_ = 5 simulating synaptic currents seen *in vivo* (Destexhe, Rudolph, Fellous, & Sejnowski, 2001):

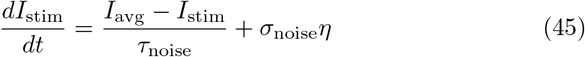

where 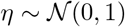, *I*_avg_ = 50*μ*A/cm^2^ and *σ*_noise_ = 25*μ*A/cm^2^. All simulations use a second-order Euler method (Moye & Diekman, 2018) to integrate the dynamics (2) for 1500 ms with a timestep of *dt* = 0.01 ms, lowered from the sampling period *T* in the interest of algorithmic stability.

**Fig. 1.**
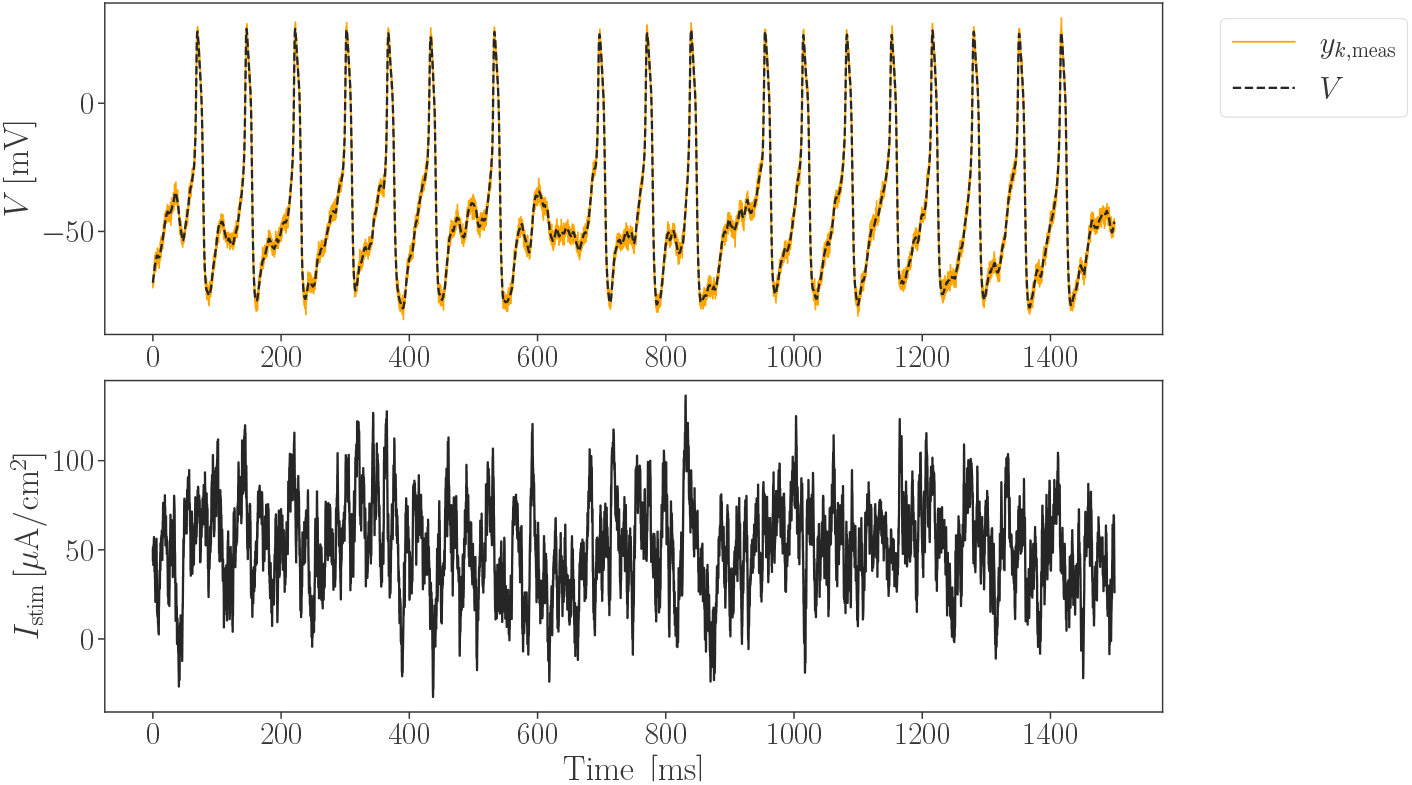
Noisy membrane voltage measurements 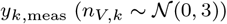 of a 2-dimensional Morris-Lecar variant (A1) (with Class 2 parameter values found in Table A) subject to a noisy input current *I*_stim_ (OU process with *I*_avg_ = 50μA/cm^2^, *τ*_noise_ = 5 and *σ*_noise_ = 25*μ*A/cm^2^)

In addition to qualitative assessments of tracking performance, errors are measured quantitatively using the root-mean-square error (RMSE).

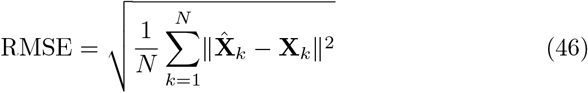

where **X**_*k*_ represents groundtruth data.

### 3.1 Exploring the filter parameter space

In performing joint state estimation and parameter identification for a conductance-based neuron model, the accuracy of the tracking and convergence to the true states are most important. Figure 2 and Figure 3 show the root-mean-square tracking error (RMSE) result of sweeps over parameters *λ*_0_, *δ*_0_ and *a, b* from equations (43) and (44) respectively. The RMSE is computed from the simulation half-point to allow enough time for transients to subside and subsequently divided by the range of each variable (see Table 1) to facilitate the comparison of tracking errors.

**Fig. 2.**
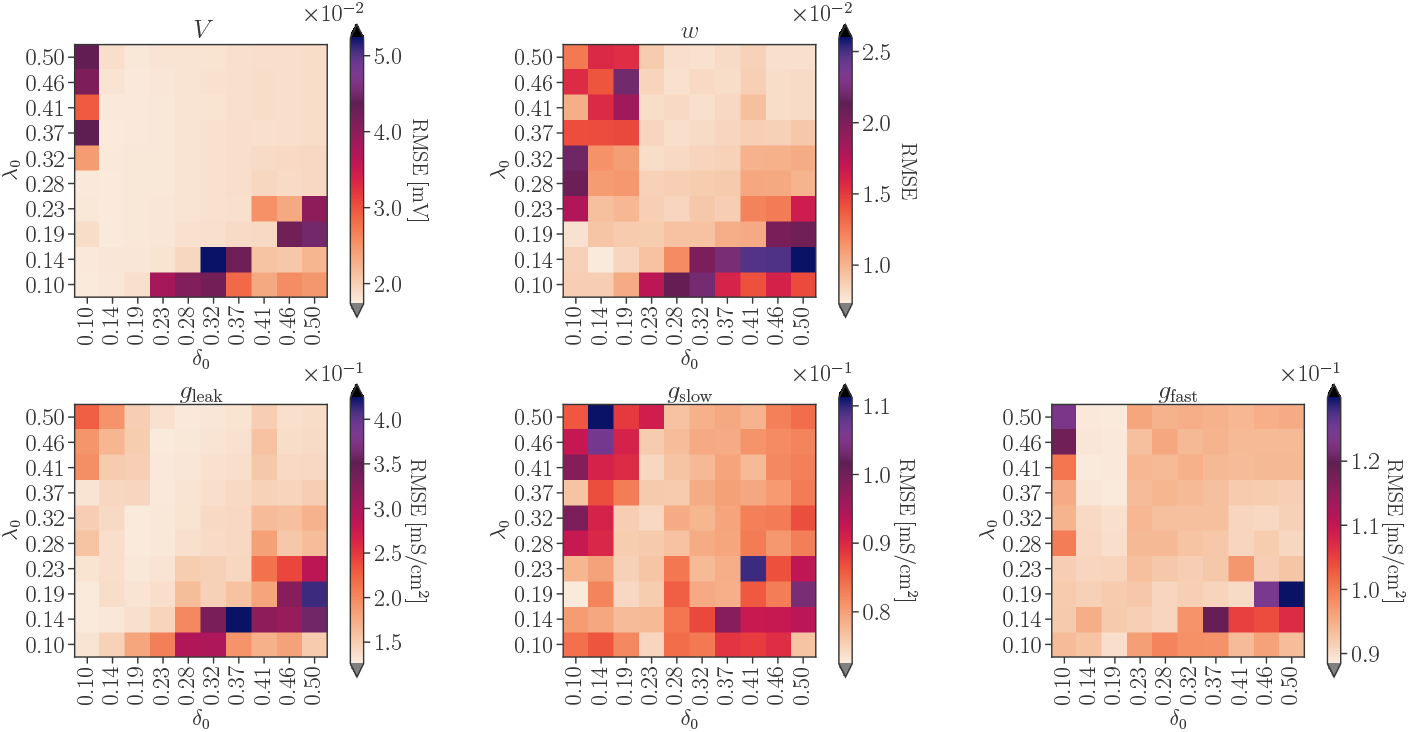
Parameter sweep over *λ*_0_ and *δ*_0_ according to RMSE (default parameter values: *a* = 5, *b* = 5, *σ* = 0.5)

**Fig. 3.**
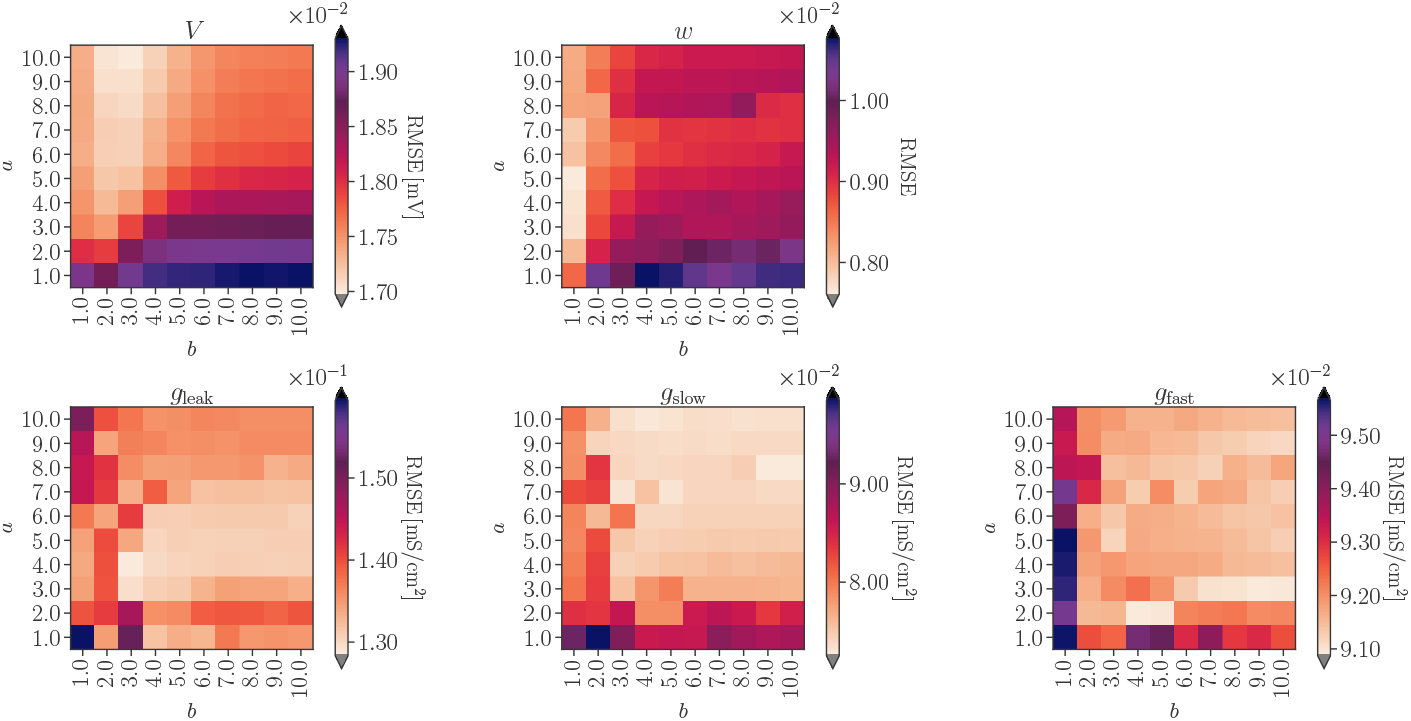
Parameter sweep over *a* and b according to RMSE (default parameter values: *λ*_0_ = 0.2, *δ*_0_ = 0.2, *σ* = 0.5)

According to Figure 2, the combination of a small *δ*_0_ and small *λ*_0_ seem to be the most effective. Yet, for larger *δ*_0_, *λ*_0_ values, the filter becomes too sensitive to adaptation leading to more frequent failures. RMSE increases when *δ*_0_ and *λ*_0_ are dissimilar, when one is much larger than the other.

The higher the *a* and *b*, the higher the probability that *λ* ← *λ*_0_ and *δ* ← *δ*_0_ respectively. While this can be seen above for *a* ≥ 5, the selection of *b* does not seem particularly sensitive. Overall, *λ*_0_ = *δ*_0_ = 0.2, *a* ≥ 5.0 and *b* ∈ [3, 10] seem appropriate for this system.

### 3.2 Joint State and Parameter Estimation

In the absence of adaptation, poor initialization of noise covariance matrices **Q** and **R** significantly impacts the stability of a recursive filter. Unless specified otherwise, the following parameterization is used throughout this section to illustrate this point: **X**_0_ = [-100, 0.5, 10, 80, 140], **P**_0_ = diag([0.0001, 0.0001, 0.0001, 0.0001]), **Q**_0_ = diag([10,0.001, 10, 10, 10]), **R**_1_ = 0.3.

Figure 4 compares the performance of the UKF algorithm and that of the RAUKF on a tracking task where the initialization of the filters is identical in **X**_0_, **P**_0_, **Q**_0_ and **R**_1_. While both filters successfully maximize the probability of observing the measurements (top panel), the UKF struggles to estimate the unobserved state (bottom panel). This shortcoming of the standard filter is even more pronounced during the identification of conductance parameters, as shown in Figure 5.

**Fig. 4.**
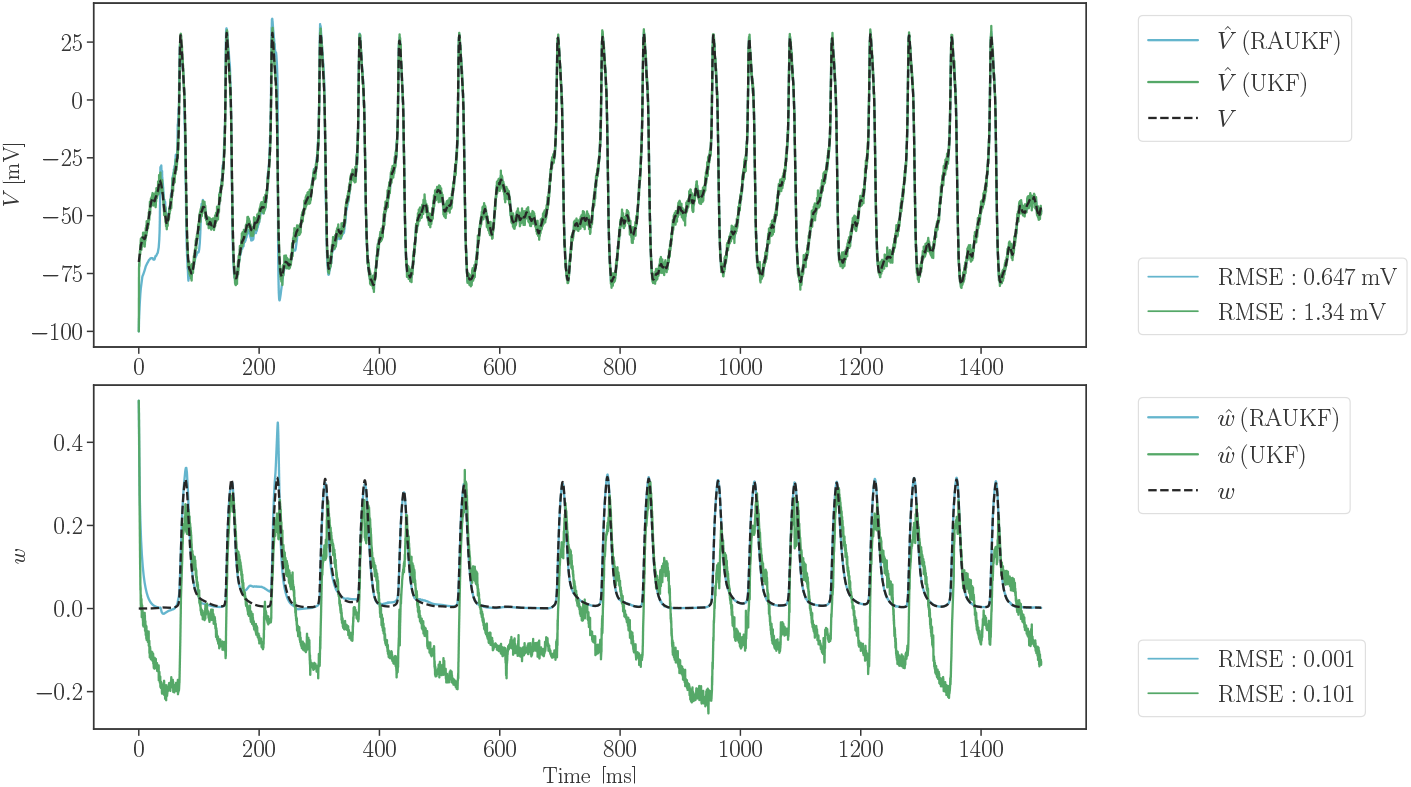
Tracking the neuron model subject to a stochastic input current *I*_stim_, noisy measurements *y* and unknown conductances

**Fig. 5.**
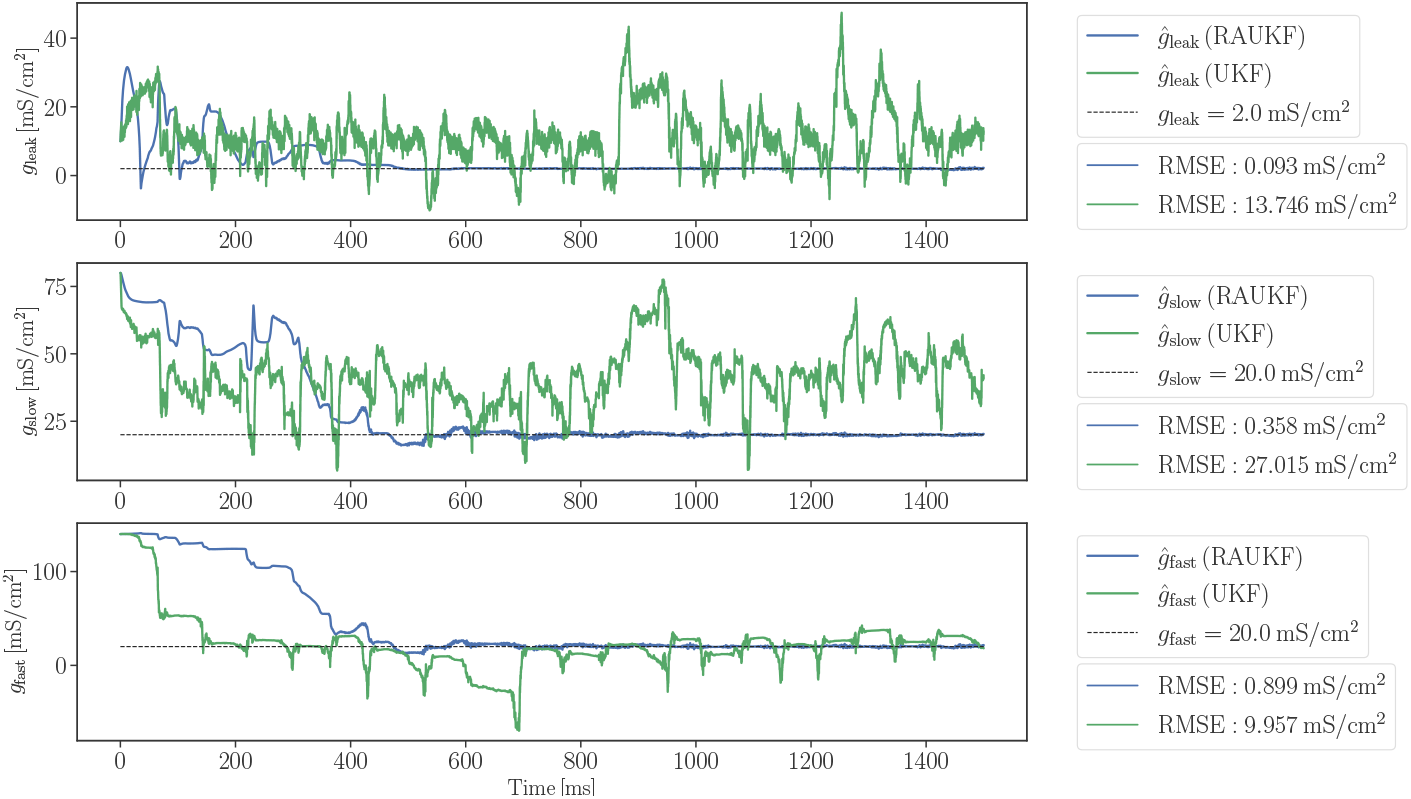
Identifying the neuron model parameters from noisy observations

In both cases, the RAUKF leverages the fault detection process to adapt the unknown covariance matrices, resulting in clear tracking and identification improvements. Transient effects of adaptation are noticeable early on in the simulation, when a high number of corrections are made (i.e., when the innovation vector is the least normally distributed). Quantitative performance of the filters is compared based on RMSEs, evaluated from *t* = 750 ms onward to minimize the impact of transients.

### 3.3 Performance Against Measurement Faults

The fault detection test introduced in section 2.5 was originally developed as a response to potential actuator or sensor malfunction. To emulate a faulty sensor, we consider a scenario where the measurement noise profile of the membrane voltage observations changes mid-simulation. In such a scenario, standard recursive estimation techniques (e.g. UKF) tend to fail owing to the unexpected change in noise covariance properties and the lack of adaptation (see the red line in Figure 6). On the other hand, as the faulty measurements momentarily alter the distribution of the innovation vector, the RAUKF triggers a correction of the measurement noise covariance matrix **R**_*k*_. Figure 6–7 illustrate the changing measurement noise profile and its effect on state tracking and parameter identification.

**Fig. 6.**
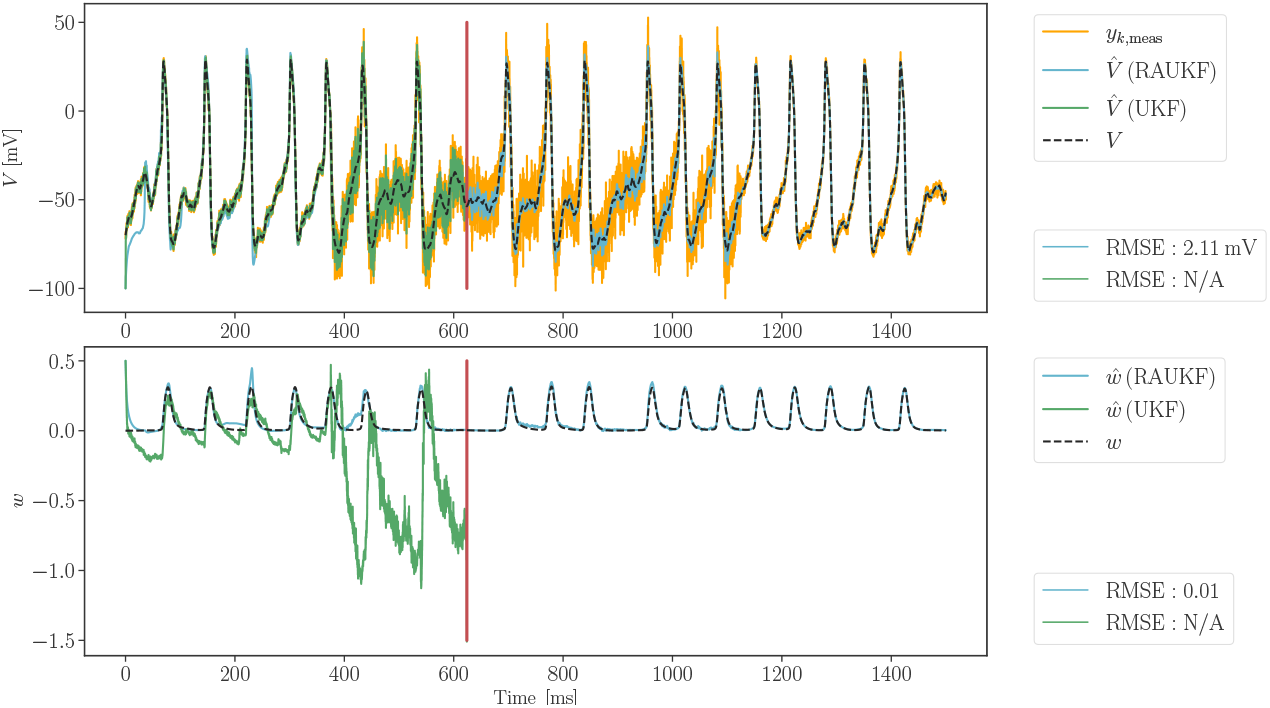
Tracking performance under momentary change in measurement noise profile to simulate faulty membrane voltage sensing (*ñ_V,t_* = 5*n_V,t_*, *t* ∈ [375,1125] ms)

**Fig. 7.**
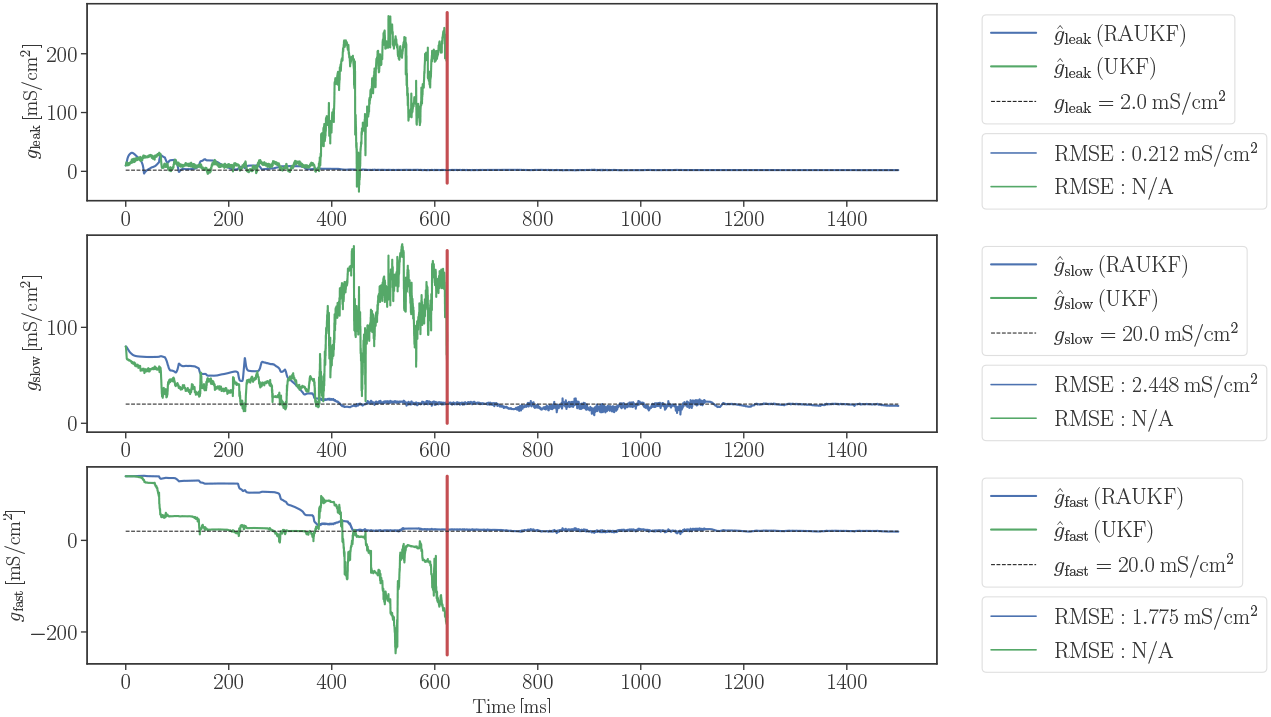
Parameter identification performance under momentary change in measurement noise profile to simulate faulty membrane voltage sensing (*ñ_V,t_* = 5*n_V,t_*, *t* ∈ [375, 1125] ms)

### 3.4 Performance Against Model Inadequacies

The final simulation tests the RAUKF tracking performance when using incomplete models. A 3-dimensional Morris-Lecar-like model (see Morris-Lecar variant (A2) with parameter values A2) is used to generate noisy observations, while the filter’s dynamics model is described by a 2-dimensional Morris-Lecar-like model (A1). As a result, the source of the measurements and the tracking model no longer match. This setup aims to simulate a practical setting where measurements of membrane voltage are far more expressive than that which could be reproduced with a low-dimensional model.

The 3-dimensional model (A2) splits the slow current *I*_slow_ = *ḡ*_slow_*w*(*V* – *E*_K_) into a K^+^ rectifier current *I_K, dr_* and a subthreshold (outward) current *I*_sub_ (Prescott et al., 2008). Since (A2) includes all the parameters from (A1), the estimation objectives remain the same as in the previous simulations, with only the measurements being different. Figure 8 and Figure 9 showcase the tracking performance achieved with RAUKF despite the model mismatch.

**Fig. 8.**
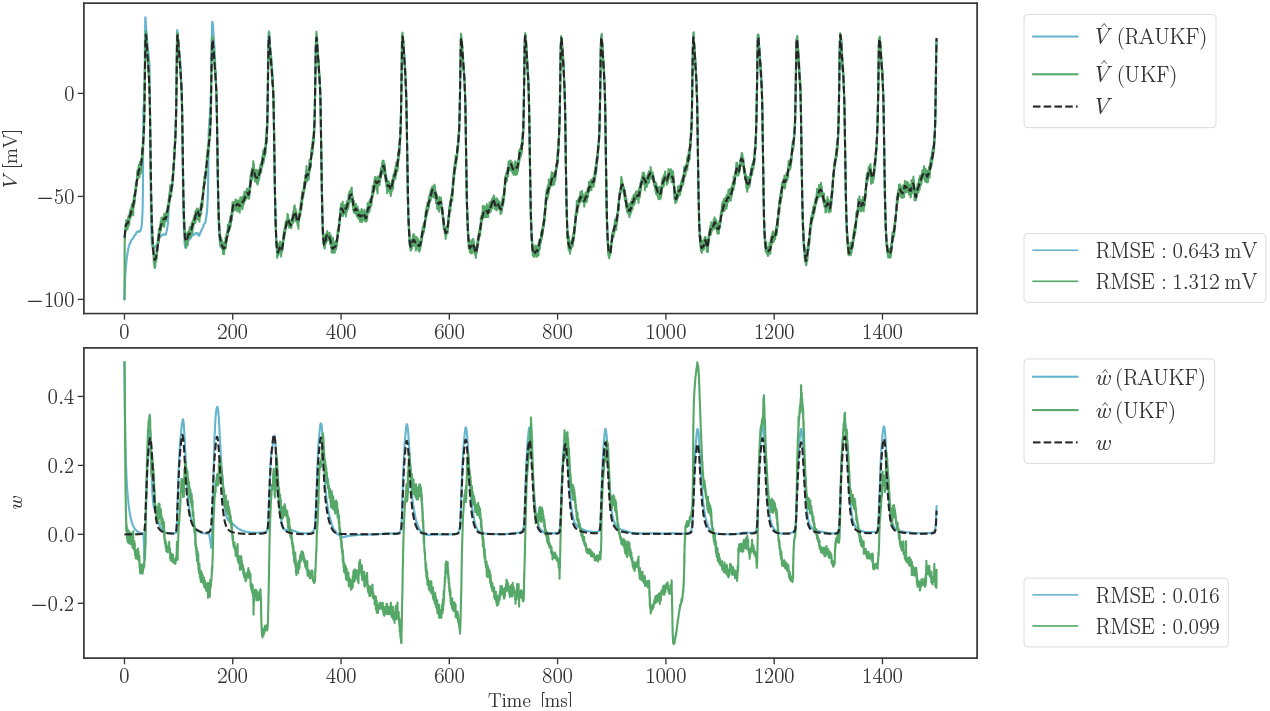
Tracking performance given a mismatch between the dynamics model and the model generating observations

**Fig. 9.**
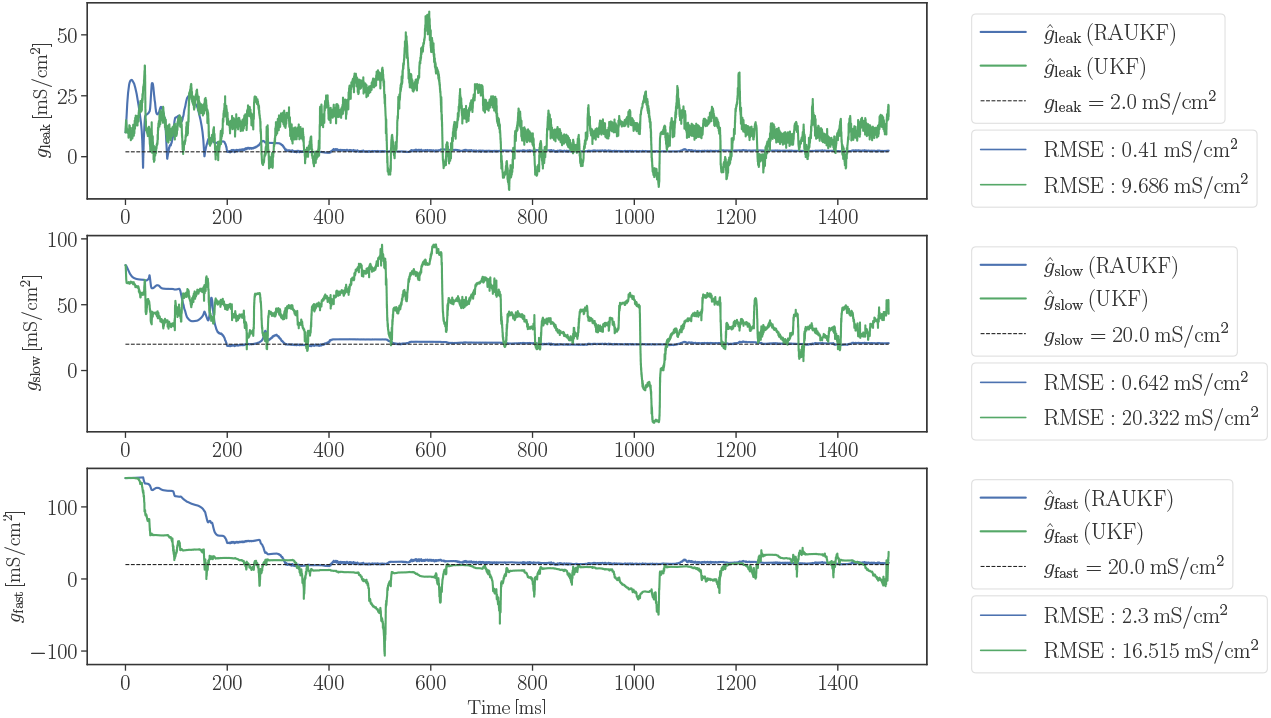
Parameter identification performance given a mismatch between the dynamics model and the model generating observations

While it comes as no surprise that the filter now favours observations over its internal estimate of the membrane voltage, the successful tracking of the unobserved state and parameters reflects the robustness of the filter. Compared to previous simulations, the estimation incurs a larger overall tracking error, yet the accuracy remains satisfactory (compare RMSEs between Figures 4–5 and Figures 8–9). The ability to accurately determine the true channel conductances from more realistic measurements is of particular importance for practical applications of these estimation methods.

## 4 Discussion

The application of state estimation in neuroscience and biomedical fields is burgeoning. In this work, we detail the efficacy of a robust and adaptive unscented Kalman filter (RAUKF) when applied to neuronal state and parameter estimation. RAUKF is capable of estimating neuronal dynamics accurately in simulation despite initial poor estimates. Further, RAUKF maintains tracking performance when subject to measurement faults and model mismatch.

Tuning parameters of the RAUKF is something that must be done under the consideration of the system that is being modelled. While some generalizations about parameter selection can be made, most are specific to the context in which the filter is being applied. The choice of parameters *a* and *b* depend on the model being used. In a previous work (Zheng et al., 2018), *a* and b were increased in tandem resulting in an increase of performance that plateaued. However, as can be seen in Figure 3 the performance along the line where *a* = *b* remains roughly the same, instead, performance increases with an increase in *a* and degrades with an increase in *b*. The performance trend observed with an increase in *a* is likely due to the filter’s poor initialization of estimates for the values and covariances of the process of which *a* has direct influence over the adaptation of (43). In contrast, the filter’s observation comes from the membrane potential, and this source is known to have a white Gaussian noise, increasing *b* makes the filter over-tune the estimate of the noise of the observation, detracting from the filter’s capacity to estimate the hidden states of the system. This difference is illustrated in the heatmap of the RMSE for the *w* gating parameter, where an increase in *b* does not lead to as drastic a decrease in performance compared to the states that are estimated by a random walk. The values to use for *λ*_0_ and *δ*_0_ may be adjusted in a similar fashion to *a* and b, as the model has less certainty about the process dynamics then the observation so there is a bias to the value of *λ*_0_ over *δ*_0_.

Fault detection is especially of use when modelling mechanisms that are prone to discontinuous or abrupt changes, such as spike firing in neurons, especially when being acted upon by some external activity. Action potentials and their related spike times are often of particular relevance when replicating a neuron’s behaviour in a model. A fault detection mechanism would make it such that spikes that are not reflected in the model’s prediction will likely be detected as a fault, resulting in a non-normal distribution such that the state estimation, more so the variance of the estimation, will change more rapidly. While not considered in this work, modifying the filter to constrain the parameter space would allow for better parameter fitting (Simon, 2010). Constraining parameter states based on their physical/biological interpretation may allow for better tracking, constraining the gating parameter *w* between [0, 1] as the value represents how open a given channel is. This type of constraint behaviour may also make the filter more robust as the parameter space being constrained would be able to prevent numerical instabilities that unconstrained estimates may tend towards.

The application of the RAUKF on incomplete models relates to possible future research that may extend this work. When the state estimation process does not match the model used to generate the observed state (Figure 8) the filter still does well to track state estimates even when the mismatch between the observation and process would be apparent. Future work may address this by introducing some of the constrained methods previously mentioned (Simon, 2010) and incorporating additional states following random walk dynamics that may be representative of hidden dynamics not encapsulated by the process model. In addition, building upon previous work conducted on simpler neuronal models (e.g., leaky integrate-and-fire) (Lankarany, Heiss, Lampl, & Toyoizumi, 2016; Lankarany, Zhu, Swamy, & Toyoizumi, 2013), robust and adaptive filtering techniques may be used to identify heterogeneous populations of neurons and possible differential properties or mechanisms (such as excitatory/inhibitory afferents or conductances) that contribute to this heterogeneity.

A possible extension would be to determine channel conductance properties of specific cell types in different locations, such as hyperpolarization-activated cation (h-) channels in oriens lacunosum-moleculare (OLM) cells of the hippocampus. H-channels in OLM cells are known to vary in a location-dependent fashion (Hilscher et al., 2019), and OLM cells are known to be important contributors of theta rhythms that facilitate spatial memory processes (Klausberger & Somogyi, 2008). We have developed detailed multi-compartment OLM cell models to understand how they contribute to circuit function (Sekulić et al., 2020). Since we have shown that the RAUKF technique can deal with incomplete models, it may be possible to determine varying h-channel conductances using reduced (and thus incomplete) OLM cell models with experimental OLM cell recordings from different locations.

Finally, it is important to note that the adaptation of noise covariance matrices presented in this work applies to a wide range of recursive state estimation algorithms. While the UKF generally performs better than other filters in its class, its sampling-based approach and higher computational cost may hinder its adoption in embedded biomedical application. Alternatives based on linearization include a robust adaptive EKF or, if the hardware supports it, a robust adaptive iterated EKF. Demonstrations of these alternatives for the joint state and parameter estimation of various neuronal models can be found in the supplementary material.

The assumption of linearizable dynamics may prove unrealistic for certain neuronal models, especially as higher-order dynamics are considered. However, extending early works on single model tracking and control (Ullah & Schiff, 2009), we believe estimation algorithms such as RAUKF could be effectively combined with low-order representations of high dimensional neuronal dynamics (e.g., reduced order models, mean-field models) in future model-based closed-loop neuroscience applications.

## Supplementary information

The Python code used in this study can be found at https://github.com/nsbspl/RAUKF.

## Appendix A Conductance-based models

The conductance-based models used in this study are derived from the Morris-Lecar model (Morris & Lecar, 1981) as shown in Prescott et al. (2008). The 2-dimensional model is given by (A1) based on the parameters in Table A.

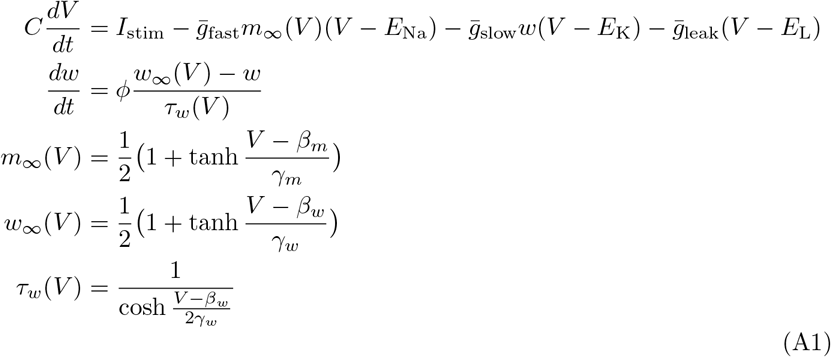

The 3-dimensional model used in the model mismatch simulation (see Figure 3.4) is given by (A2) based on the parameters in Table A2.

**Table A1.** Parameters for the 2-dimensional Class 1 (2) [3] Morris-Lecar-like model

| Parameter | Value | Units |
| --- | --- | --- |
| $C$ | 2 | $\mu\text{Fcm}^{-2}$ |
| $\phi_w$ | 0.15 | $\text{s}^{-1}$ |
| $\bar{g}_{\text{fast}}$ | 20 | $\text{mS cm}^{-2}$ |
| $E_{\text{Na}}$ | 50 | mV |
| $\bar{g}_{\text{slow}}$ | 20 | $\text{mS cm}^{-2}$ |
| $E_{\text{K}}$ | -100 | mV |
| $\bar{g}_{\text{leak}}$ | 2 | $\text{mS cm}^{-2}$ |
| $E_{\text{L}}$ | -70 | mV |
| $\beta_m$ | -1.2 | mV |
| $\gamma_m$ | 18 | mV |
| $\beta_w$ | 0 (-13) [-21] | mV |
| $\gamma_w$ | 10 | mV |

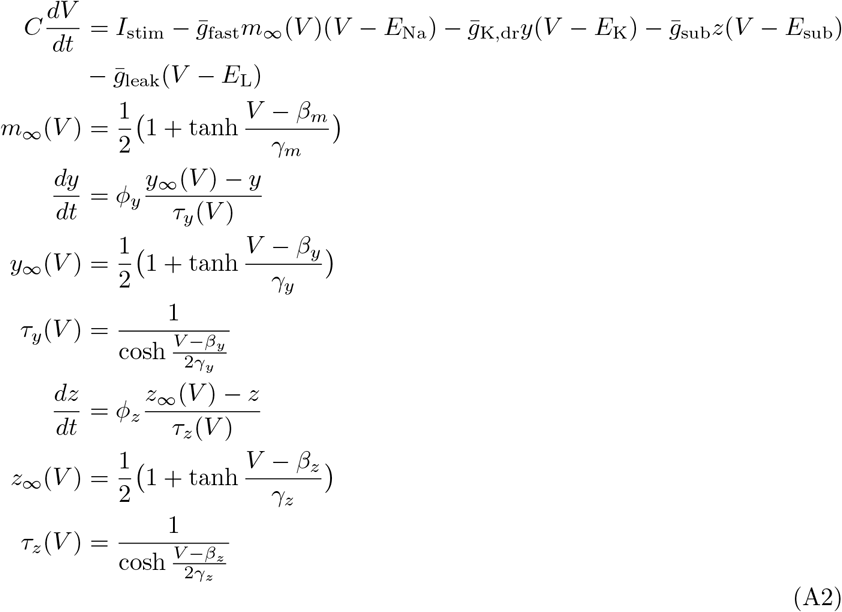

**Table A2.**
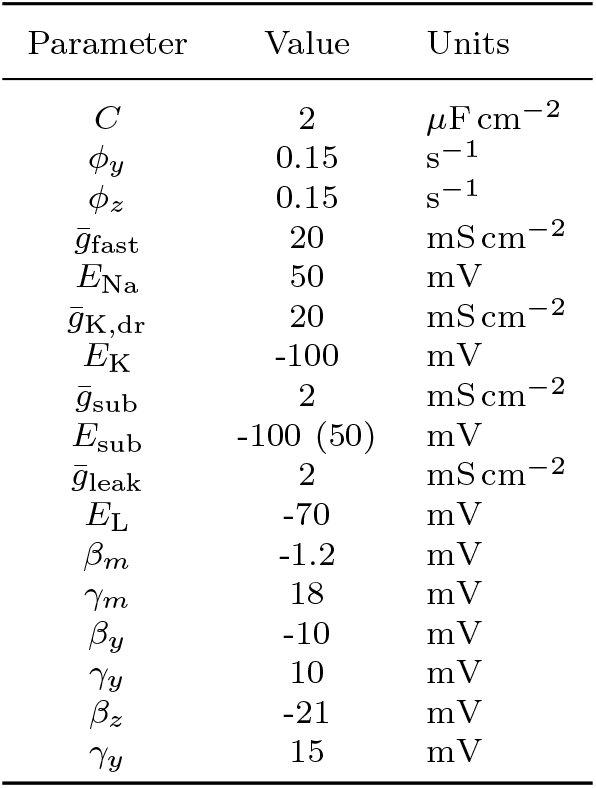
Parameters for the 3-dimensional Morris-Lecar-like model with outward (inward) *I*_sub_

## References

Barfoot, T.D. (2017). State estimation for robotics. Cambridge University Press.

Destexhe, A., Rudolph, M., Fellous, J.M., Sejnowski, T.J. (2001). Fluctuating synaptic conductances recreate in vivo-like activity in neocortical neurons. Neuroscience, 107(1), 13–24.

Golowasch, J. (2014). Ionic current variability and functional stability in the nervous system. BioScience, 64 (7), 570–580.

Hajiyev, C., & Caliskan, F. (2003). Fault diagnosis and reconfiguration in flight control systems. Boston: Kluwer Academic Publishers.

Hajiyev, C., & Soken, H.E. (2014). Robust adaptive unscented kalman filter for attitude estimation of pico satellites. International Journal of Adaptive Control and Signal Processing, 28(2), 107–120.

Hilscher, M.M., Nogueira, I., Mikulovic, S., Kullander, K., Leão, R.N., Leão, K.E. (2019). Chrna2-olm interneurons display different membrane properties and h-current magnitude depending on dorsoventral location. Hippocampus, 29(12), 1224–1237.

Julier, S., & Uhlmann, J. (1997). New extension of the Kalman filter to nonlinear systems. I. Kadar (Ed.), Signal processing, sensor fusion, and target recognition vi (Vol. 3068, pp. 182 – 193). SPIE.

Julier, S., Uhlmann, J., Durrant-Whyte, H. (2000). A new method for the nonlinear transformation of means and covariances in filters and estimators. IEEE Transactions on Automatic Control, 45(3), 477–482.

Kalman, R.E. (1960). A New Approach to Linear Filtering and Prediction Problems. Journal of Basic Engineering, 82(1), 35–45.

Klausberger, T., & Somogyi, P. (2008). Neuronal diversity and temporal dynamics: The unity of hippocampal circuit operations. Science, 321 (5885), 53–57.

Lankarany, M., Heiss, J.E., Lampl, I., Toyoizumi, T. (2016). Simultaneous bayesian estimation of excitatory and inhibitory synaptic conductances by exploiting multiple recorded trials. Frontiers in Computational Neuroscience, 10(nil), nil.

Lankarany, M., Zhu, W.-P., Swamy, M. (2014). Joint estimation of states and parameters of hodgkin-huxley neuronal model using kalman filtering. Neurocomputing, 136(nil), 289–299.

Lankarany, M., Zhu, W.-P., Swamy, M.N.S., Toyoizumi, T. (2013). Inferring trial-to-trial excitatory and inhibitory synaptic inputs from membrane potential using gaussian mixture kalman filtering. Frontiers in Computational Neuroscience, 7(nil), nil.

Mohamed, A.H., & Schwarz, K.P. (1999). Adaptive kalman filtering for ins/gps. Journal of Geodesy, 73(4), 193–203.

Morris, C., & Lecar, H. (1981). Voltage oscillations in the barnacle giant muscle fiber. Biophysical journal, 35(1), 193–213.

Moye, M.J., & Diekman, C.O. (2018). Data assimilation methods for neuronal state and parameter estimation. The Journal of Mathematical Neuroscience, 8, 11.

Prescott, S.A., De Koninck, Y., Sejnowski, T.J. (2008). Biophysical basis for three distinct dynamical mechanisms of action potential initiation. PLOS Computational Biology, 4 (10), 1–18.

Schiff, S.J. (2009). Kalman meets neuron: the emerging intersection of control theory with neuroscience. Annual International Conference of the IEEE Engineering in Medicine and Biology Society. IEEE Engineering in Medicine and Biology Society. Annual International Conference, 2009, 3318–3321.

Schiff, S.J. (2011). Neural Control Engineering: The Emerging Intersection between Control Theory and Neuroscience. The MIT Press.

Sekulic, V., Yi, F., Garrett, T., Guet-McCreight, A., Lawrence, J.J., Skinner, F.K. (2020, sep). Integration of within-cell experimental data with multicompartmental modeling predicts h-channel densities and distributions in hippocampal OLM cells. Frontiers in Cellular Neuroscience, 14.

Simon, D. (2010). Kalman filtering with state constraints: A survey of linear and nonlinear algorithms. IET Control Theory and Applications, 4 (8), 1303–1318.

Skinner, F. (2006). Conductance-based models. Scholarpedia, 1 (11), 1408.

Stengel, R.F. (1994). Optimal control and estimation. Dover Publications.

Ullah, G., & Schiff, S.J. (2009). Tracking and control of neuronal hodgkin-huxley dynamics. Phys. Rev. E, 79, 040901.

Voss, H., Timmer, J., Kurths, J. (2004). Nonlinear dynamical system identification from uncertain and indirect measurements. Int. J. Bifurc. Chaos, 14, 1905–1933.

Zheng, B., Fu, P., Li, B., Yuan, X. (2018). A robust adaptive unscented kalman filter for nonlinear estimation with uncertain noise covariance. Sensors, 18(3), 808.

